# Modeling the MYC-driven normal-to-tumour switch in breast cancer

**DOI:** 10.1101/380931

**Authors:** Corey Lourenco, Manpreet Kalkat, Kathleen E. Houlahan, Jason De Melo, Joseph Longo, Susan J. Done, Paul C. Boutros, Linda Z. Penn

## Abstract

The potent MYC oncoprotein is deregulated in many human cancers, including breast carcinoma, and is associated with aggressive disease. To understand the mechanisms and vulnerabilities of MYC-driven breast cancer, we have generated an *in vivo* model that mimics human disease in response to MYC deregulation. MCF10A cells ectopically expressing a common breast cancer mutation in the PI3 kinase pathway (PIK3CA^H1047R^) lead to the development of organized acinar structures in mice. However, expressing both PIK3CA^H1047R^ and deregulated-MYC lead to the development of invasive ductal carcinoma, thus creating a model in which a MYC-dependent normal-to-tumour switch occurs *in vivo*. These MYC-driven tumors exhibit classic hallmarks of human breast cancer at both the pathological and molecular levels. Moreover, tumour growth is dependent upon sustained deregulated MYC expression, further demonstrating addiction to this potent oncogene and regulator of gene transcription. We therefore provide a MYC-dependent human model of breast cancer which can be assayed for *in vivo* tumour initiation, proliferation, and transformation from normal breast acini into invasive breast carcinoma. Taken together, we anticipate that this novel MYC-driven transformation model will be a useful research tool to both better understand MYC’s oncogenic function and identify therapeutic vulnerabilities.

**Conflict of interest statement:** The authors declare no potential conflicts of interest.

## INTRODUCTION

MYC deregulation occurs in the majority of human cancers and is therefore an ideal therapeutic target.^1^ Deregulation can be broadly defined as any genomic, epigenetic or signaling pathway aberration that results in sustained and often elevated MYC expression and activity. However, targeting this potent oncoprotein has been difficult using traditional approaches as MYC is a transcription factor lacking enzymatic pockets. Alternative strategies are therefore under intense investigation, including targeting MYC:DNA interaction with small protein inhibitors such as Omomyc or ME47.^2,3^ In addition, identifying the unique vulnerabilities which arise in a MYC-driven tumour can reveal targetable synthetic lethal interactions.^4–8^ There is therefore a need for a diverse set of MYC-driven models to identify and test new therapeutic strategies in a MYC-transformed setting.

Many approaches have been developed to model MYC in cancer, including cancer cell lines and genetically engineered mouse models (GEMMs). Rodent models include the widely used Rat-1A cell line, the Eμ-MYC transgenic model of B-cell lymphoma and the MMTV-MYC model of breast cancer (BC).^9–12^ Additionally, human MYC-dependent models, such as the Burkitt lymphoma cell line model P493-6 with regulatable MYC levels, have been used extensively to interrogate transcriptional regulation by MYC.^13–15^ Moreover, the MycER chimeric protein system, which can be applied to cell line and mouse models of many cancer types, has been used for studies of MYC in the cell cycle, genomic instability, transformation and synthetic lethal interactions.^16–22^ These models have provided several new insights into MYC biology, suggesting further development of diverse models will also provide important contributions, particularly if they are complementary and obviate associated limitations with existing models. For instance, GEMMs are time-consuming to manipulate genetically, are resource-intensive and often do not accurately model all human disease at the histological level. Specifically, when modeling human breast cancer, there are numerous differences between mouse and human mammary gland development and the surrounding stroma that need to be considered.^23^ In addition, most cancer cell line models do not accurately reflect human tumours *in vivo*, are derived from already fully transformed tumour tissue, and are not necessarily driven by or dependent on deregulated MYC. We have therefore set out to design a functionally complementary MYC-driven transformation model that meets the following criteria: is fully-transformed in response to, and dependent-upon, deregulated MYC; recapitulates human disease at both the pathological and molecular levels *in vivo*; and, is easy to manipulate genetically. These criteria outline a model that best reflects the complexity of human tumours *in vivo* while ensuring that identifying MYC-dependencies is possible. To this end, we used a non-transformed parental cell (MCF10A) and developed an isogenic panel of cell lines that could meet these standards and model the transformation of BC upon the introduction of deregulated MYC. We evaluate MYC-dependency in vivo and characterize the histology and mRNA expression changes that occur in the these MYC-driven tumours.

## RESULTS and DISCUSSION

### *In vivo* transformation of MCF10A cells is dependent on active PI3K signaling, deregulated MYC, and requires MBII

We chose to model BC as MYC is often deregulated and contributes to the progression of this disease.^24,25^ Recent large whole-genome sequencing studies of BC patient tumours revealed that the most frequently mutated and amplified oncogenes in BC are PIK3CA and MYC, respectively.^26^ PIK3CA is mutated in approximately 30% of BC patient tumours, whereas MYC deregulation, primarily scored through amplification, occurs in approximately 20% of BCs.^26,27^ To model BC transformation, we sequentially introduced these commonly mutated oncogenes into the MCF10As, an immortal, non-transformed breast cell line.^28^ MCF10As were chosen due to their inherently stable genome, which ensures the oncogenic conversion into the transformed state is due to the introduction of ectopic oncogenes rather than an indirect consequence of genomic instability and the selection of a transformed clone. In addition, the MCF10As are a relevant model as the few mutations acquired during spontaneous immortalization are prevalent in human BC, including CDKN2A deletion.^29^ Finally, despite harbouring a focal amplification at chromosome 8q24, total MYC levels remain highly regulated, decreasing rapidly in response to anti-proliferative conditions which is a hallmark of normal MYC regulation in non-transformed cells.^29,30^ Thus, the MCF10A cells fulfilled our criteria of a parental breast cell system to base our model.

We first stably introduced the empty vector control or the activated PIK3CA^H1047R^ allele into MCF10A cells. The latter was active in these cells as AKT phosphorylation at serine 473 was readily detectable (Figure 1A). To this pair of cell lines, we introduced empty vector control or ectopic constitutive MYC expression, to model MYC deregulation. This resulted in a panel of four cell lines consisting of 10A.EE (empty vector, empty vector), 10A.EM (empty vector, MYC), 10A.PE (PIK3CA, empty vector) and 10A.PM (PIK3CA, MYC; Figure 1A). Under growing conditions, we observed no significant change in either cell size (Supplementary Figure 1A) or proliferation (Figure 1B) between any groups, suggesting that these cellular properties would not contribute to any phenotypes observed. Additionally, despite minor cell morphology differences (Supplementary Figure S1B), epithelial-to-mesenchymal (EMT) markers (Supplementary Figure S1C) did not significantly change after activation of the PI3K pathway.

**Figure 1.**
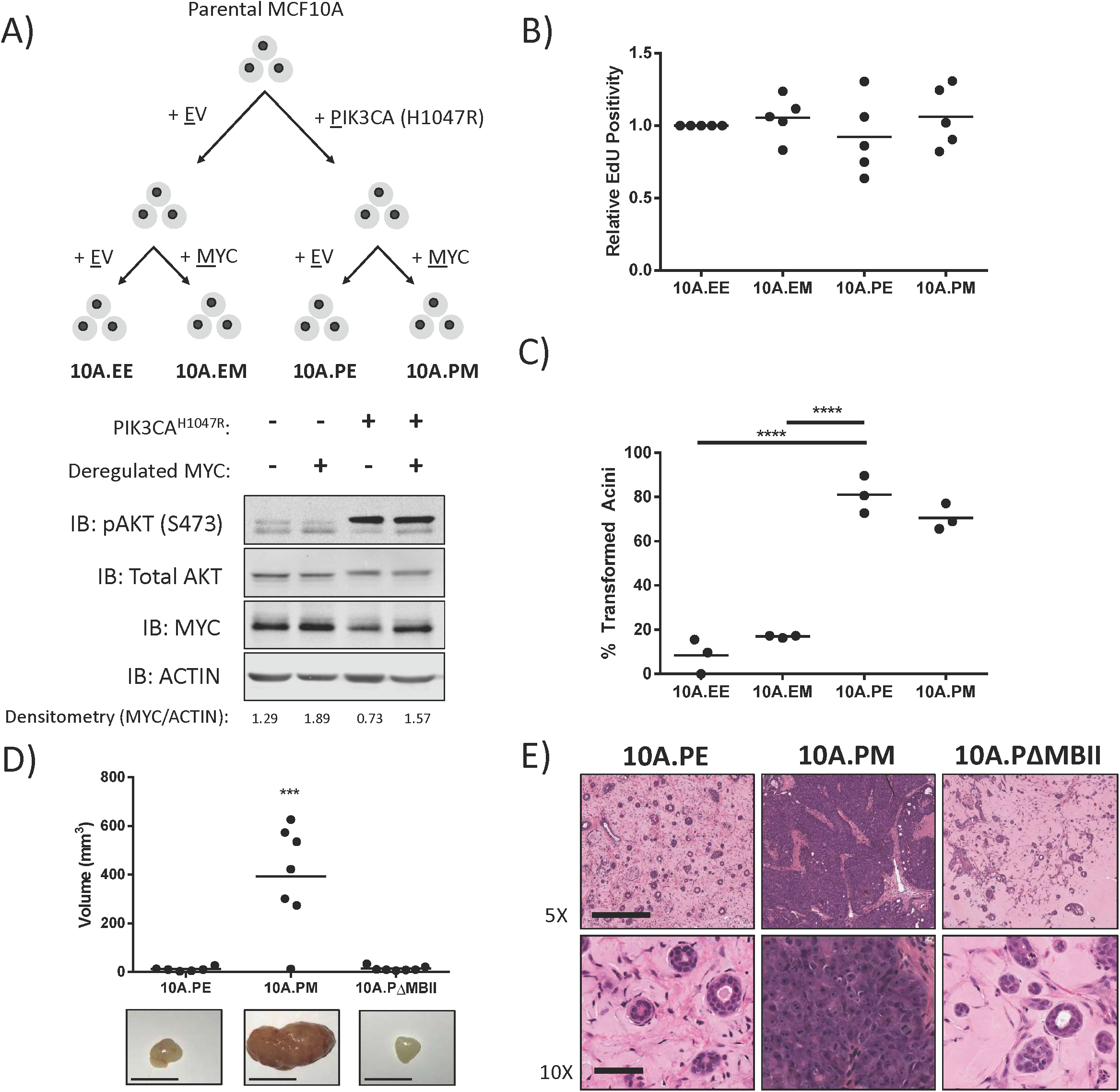
*In vivo* transformation of MCF10A is dependent on active PI3K signaling, deregulated MYC and requires MBII. **(A)** Schematic overview for the generation of an isogenic panel of MCF10A cell lines. Immunoblotting was performed against pAKTS473, total AKT, MYC and Actin. Quantification of MYC and ACTIN intensity done with ImageJ. **(B)** EdU proliferation measurements for the isogenic MCF10A panel. Mean values are shown, one-way ANOVA with Bonferroni post-test for multiple testing, 5 biological replicates. **(C)** The MCF10A isogenic panel cultured in Matrigel for 12 days. Images were acquired on day 12 and quantified based on assessment of acinar circularity and size. Individual mean values from three biological replicates are shown, ****P< 0.0001, one-way ANOVA with Bonferroni post-test for multiple testing. Scale bar = 500 μm **(D)** Individual tumour volume measurements from 6x 10A.PE, 7x 10A.PM, and 7x 10A.ΔMBII xenografts 49 days after injection. Representative images are below. Scale = 0.5 cm. ***P ≤ 0.001, one-way ANOVA with Bonferroni post-test for multiple testing. **(E)** Histology samples from xenografts in **(D)** stained with hematoxylin and eosin. Representative images are shown. Scale for 5X images = 50 μm. Scale for 10X images = 500 μm.

To assay for transformation, we cultured our isogenic cell line panel as breast organoids on Matrigel to model three-dimensional acini formation *in vitro* (Figure 1C). ^31^ 10A.EE and 10A.EM cells formed round, organized acinar structures consistent with previous publications.^30^ In contrast, 10A.PE and 10A.PM cells formed large and disorganized acinar structures, demonstrating a transformed phenotype driven by PIK3CA activation *in vitro*, but not by MYC deregulation alone (Supplementary Figure S1D). Importantly, breast organoids are grown in specialized culturing conditions and cells undergo polarization programs while interacting with matrigel, whereas standard tissue culture lack these variables. These differences are likely important for the dramatic phenotypic differences observed.

Considering the importance that environmental context has on the phenotypes observed, we progressed to evaluate xenograft growth *in vivo*. This xenograft assay recapitulates the tumour microenvironment and offers an increasingly relevant context to study tumour formation and growth. To assay xenograft growth, we generated cells expressing PIK3CA^H1047R^ and then introduced empty vector (10A.PE), MYC (10A.PM) or delta MYC Box II (10A.PΔMBII) (Supplementary Figure S1E). The latter serves as a negative control as MBII is evolutionarily conserved and required for MYC-dependent transformation.^9,32^ 10A.PE cells formed small but measurable masses *in vivo*, unlike parental MCF10As and 10A.EM cells, which did not establish any mass (Supplementary Figure S1F). This suggests that activation of the PI3K signaling pathway is required for the establishment of MCF10A xenografts, reflective of the *in vitro* organoid data suggesting that these cells are more transformed. By contrast, 10A.PM cells were able to form significantly larger masses compared to 10A.PE cells (Figure 1D), which was surprising as MYC did not potentiate transformation of 10A.PE cells in our *in vitro* organoid assays (Figure 1C). Cells expressing ΔMBII (10A.PΔMBII) formed small palpable masses, similar to 10A.PE, demonstrating that tumour xenograft growth was dependent on MYC and the conserved MBII region (Figure 1D). Doubling times were 6 days for 10A.PM and 60 days for 10A.PE and 10A.PΔMBII xenografts. Histological examination showed that 10A.PE and 10A.PΔMBII xenograft masses were composed of acinar structures embedded within an extracellular matrix, strikingly similar to the histology of normal human breast acini (Figure 1E). By contrast, 10A.PM xenografts possessed major phenotypic differences, appearing as dense, fully-transformed breast tumours with features reminiscent and highly similar to human disease. While active PI3K signaling transforms breast organoids *in vitro*, both PIK3CA and deregulated MYC are required for transformation *in vivo*.

### Deregulated MYC transforms breast acini into invasive ductal carcinoma *in vivo*

In clinical practice, tumours are evaluated through, histology, immunohistochemistry (IHC) and/or molecular profiling to characterize tumour subtype and features of aggressive disease. We applied these analyses to our xenografts, in a blinded-manner, with the assistance of a practicing BC pathologist. 10A.PM and 10A.PΔMBII growths had significantly increased Ki67 staining compared to 10A.PE whereas TUNEL staining was low in all samples (Figure 2A). Interestingly, while 10A.PΔMBII tumours had an increase in Ki67 signal, the overall tumour volume was equal to 10A.PE xenograft nodules. This suggests that MYC^ΔMBII^ is able to confer some proliferative advantage due to retention of partial MYC-related activity, but is not able to drive transformation, which is consistent with previous reports.^9,32^ We also analyzed clinically relevant markers such as the estrogen and progesterone receptors (ER and PR), human epidermal growth factor receptor 2 (HER2), and cytokeratin 5 (CK5) as a marker of basal cells (Figure 2B, left). The MCF10A cell line has been previously characterized as basal and triple-negative for breast cell receptors *in vitro.*^33^ Introduction of PIK3CA^H1047R^ and MYC maintained the basal and triple-negative state *in vivo*.

**Figure 2.**
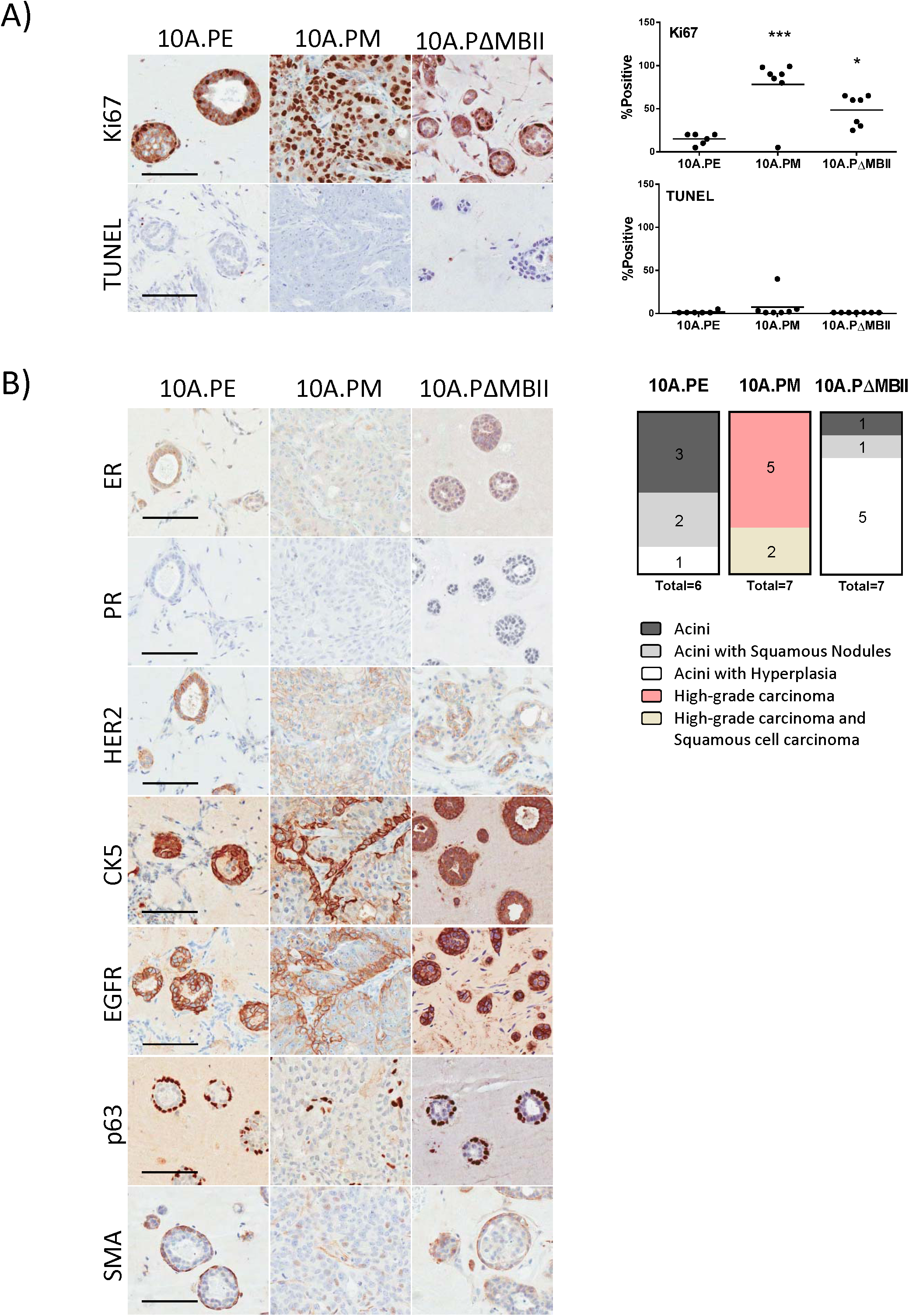
Deregulated MYC transforms breast acini into invasive breast carcinoma *in vivo.* **(A)** Tumours harvested in Figure 1C were stained (left) for Ki67 and TUNEL and were quantfied (right) for the degree of Ki67 and TUNEL staining. Individual qunatifications per tumour are shown, *P<0.05, ***P ≤ 0.001, one-way ANOVA with Bonferroni post-test for multiple testing. Scale = 100μm. **(B)** Tumours harvested in Figure 1C were stained (left) for ER, PR, HER2, CK5, EGFR, p63 and SMA by the PRP laboratory. A pathology report (right) was produced for each tumour using the combination of H&E as shown in Figure 1D and IHC markers used in this study. Scale = 100μm.

We then used clinical markers to distinguish the degree of transformation within each xenograft. Malignant disease can be categorized as either carcinoma *in situ* (CIS) or invasive ductal carcinoma (IDC) and is characterized by transformed cells contained within or invading through a layer of myoepithelial cells surrounding the acinar structure, respectively. To assess these conditions, we stained for myoepithelial cells using two markers, P63 and SMA. 10A.PE and 10A.PΔMBII acini were surrounded by myoepithelial cells and were importantly identified with human-specific epidermal growth factor receptor (EGFR) antibody, indicating that these cells were of MCF10A origin. However, 10A.PM tumours lacked p63 and SMA co-expression, suggesting a lack of myoepithelial cells in MYC-driven tumours. Our pathology report determined that a majority of 10A.PE growths were normal breast acinar structures with few cases of atypical hyperplasia (AH) (Figure 2B, right). 10A.PΔMBII growths developed moderately more AH lesions than 10A.PE xenografts, despite MBII being essential for tumour growth. These data align with our Ki67 proliferation data and previously published results showing MBII is required for transformation but can retain some MYC-related activity in tissue culture assays.^9,32^ By contrast, 10A.PM tumours, which were highly proliferative and lacked myoepithelial cells, were scored as high-grade IDC. Therefore, the addition of deregulated MYC in this *in vivo* model system transforms cells into IDC in an MBII-dependent manner. Thus, we are able to evaluate MYC-driven tumour progression (10A.PM) against a non-transformed control (10A.PE) *in vivo*, as well as measure tumour volume, and evaluate tumour pathology; a rich dataset that to the best of our knowledge uniquely distinguishes this model from previous solid human MYC-driven cancer models.

### 10A.PM tumours have MYC and invasive ductal carcinoma expression signatures

Our histological analysis demonstrates that the *in vivo* 10A.PE and 10A.PM isogenic tumour pair represents normal breast acini and IDC, respectively. We took this opportunity to identify an *in vivo* MYC-dependent transformation expression dataset. We re-established 10A.PE and 10A.PM xenografts and isolated RNA from 3 biological replicates for RNA sequencing (RNA-seq) (Figure 3A). Contaminating mouse RNA was removed using Xenome,^34^ leaving 5000 significantly differentially expressed genes (Supplementary Table S1). We performed gene set enrichment analysis (GSEA)^35^ on significant gene changes to generate a gene ontology (GO) map of up-and down-regulated biological functions in 10A.PM tumours (Figure 3B, Supplementary Table S2). Up-regulated biological functions included ribosomal biogenesis, RNA processing and translational initiation. Interestingly, down-regulated processes included cell-cell adhesion and cell differentiation, reflective of aggressive BC. Additionally, we performed GSEA to compare to well-established MYC-driven and IDC gene sets (Figure 3C, Supplementary Table S3).^36^ The MYC_UP.V1_UP gene set was significantly enriched in the up-regulated portion of our xenograft gene expression list whereas the MYC_UP.V1_DN gene list was significantly enriched in the down-regulated portion of our xenograft gene expression list. These data indicate that the expression of known up-and down-regulated MYC target genes were altered in the 10A.PM model. Additionally, we observed significant and positive NES scores with IDC gene sets such as POOLA_INVASIVE_BREAST_CANCER_UP and VANTVEER_BREAST_CANCER_POOR_PROGNOSIS indicating that these gene sets were identified in the up-regulated portion of our xenograft gene expression list. These data reinforce that our MYC-driven BC model accurately recapitulates the activated and downregulated signaling pathways associated with MYC deregulation and IDC observed in human disease.

**Figure 3.**
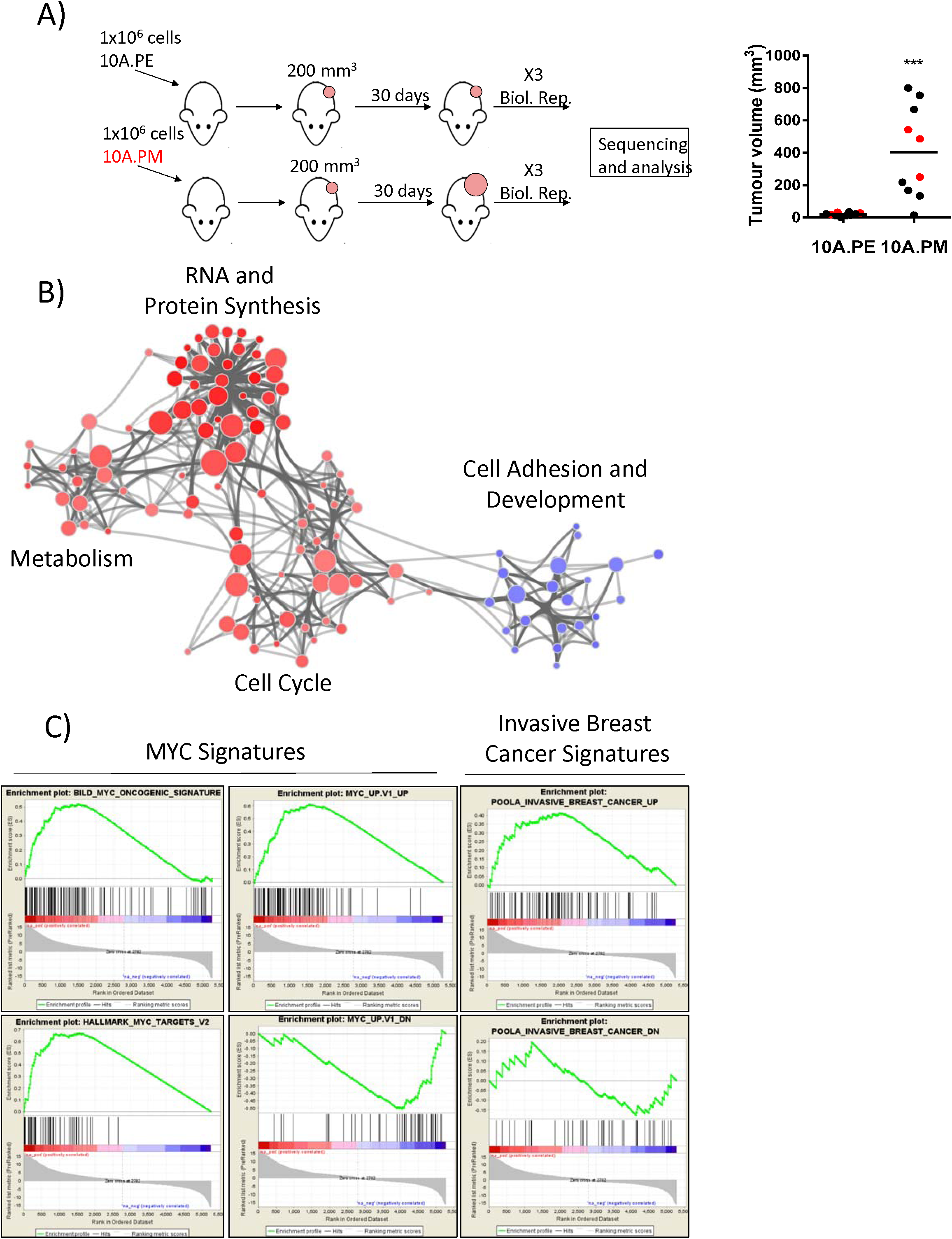
10A.PM tumours have MYC and invasive breast cancer expression signatures. **(A)** Female NOD-SCID mice were injected with 10A.PE and 10A.PM cells and allowed to form tumours (left). Three actively growing tumours (right) were harvested (red) for RNA-seq analysis. ***P<0.001, unpaired t-test. **(B)** Differentially expressed 10A.PM genes, in reference to 10A.PE genes, were mapped to display GO Biological Process gene sets (left). Edges represent the number of overlapping genes within each GO process, node size represents the number of genes within a single GO process, and node colour indicates whether a GO process contained genes which were up-regulated (red) or down-regulated (blue) in 10A.PM tumours. Darker shading indicates smaller p-value. **(C)** All differentially expressed genes were used for GSEA with MYC and IDC signature data sets. Gene sets used are labeled in figure image. Statistical analysis in supplementary table S3.

### Constitutive expression of deregulated MYC is required for tumour growth

Introducing deregulated MYC into MCF10A cells expressing PIK3CA^H1047R^ transforms these cells *in vivo*; however, it is not clear whether deregulated MYC is necessary to sustain tumour growth. To address this, we generated cell lines with doxycycline (DOX) inducible MYC to regulate ectopic expression (Figure 4A, left). Cells were treated with DOX to induce ectopic MYC expression and injected sub-cutaneously into NOD-SCID mice, which were given supplemented DOX-treated drinking water. Once tumours reached 200 mm^3^, mice were randomly assigned to ‘MYC ON’ or ‘MYC OFF’ groups, the latter being removed from DOX-supplemented drinking water (Figure 4A, right). Over 12 days of treatment, MYC ON tumours continued to grow and MYC OFF tumours slightly declined (Figure 4B, left). After 12 days of treatment, the first MYC ON tumour reached humane endpoint (Figure 4B, right). Treatment was continued for 31 days and the overall time to humane endpoint for MYC ON tumours was significantly shorter compared to MYC OFF tumours (Figure 4C). MYC ON tumours had a doubling time of 7-8 days, whereas MYC OFF tumour growth was significantly diminished (Figure 4B). This data demonstrates that this model is both driven by, and addicted to, deregulated MYC in a reproducible and quantitative manner. Furthermore, we show that targeting the deregulated fraction of MYC is sufficient to inhibit tumour growth as endogenous MYC expression is not affected by DOX treatment (Figure 4A). Therefore, the observed phenotypes, signaling pathways, and vulnerabilities that arise *in vivo* are a consequence of the regulatable and ectopic allele of MYC and importantly, are not due to permanent genomic or microenvironmental changes after deregulated MYC-driven transformation.

**Figure 4.**
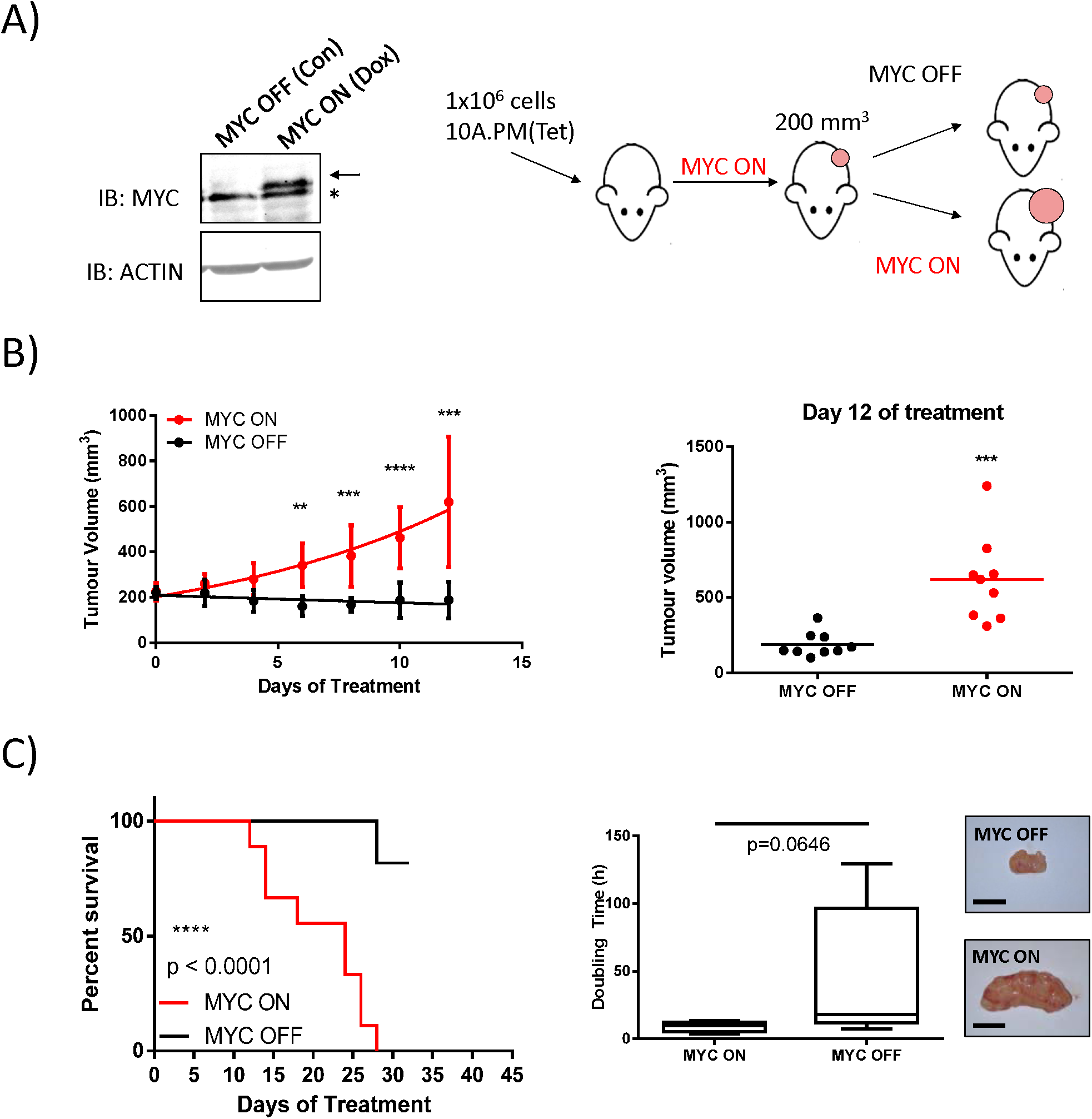
Constitutive expression of deregulated MYC is required for tumour growth. **(A)** With the addition of 100 μg/mL Doxycycline (DOX, Sigma), ectopic MYC is expressed (MYC ON). Arrow indicates ectopic MYC, * indicates endogenous MYC. 1×10^6^ cells with DOX-induced MYC expression were injected into female NOD-SCID mice and allowed to grow to 200mm^3^. Animals were then randomized into MYC ON or MYC OFF groups. **(B)** Individual mean values from all measured tumours (9 per group) during the treatment period are shown (left) and individual tumour volumes from day 12 (final day with all animals present) are also shown (right), **P<0.01, ***P<0.001, ****P< 0.0001, unpaired t-test. Scale = 0.5 cm. Error bars represent standard deviation. **(C)** Left, tumours were measured until the mice reached a humane endpoint of 1000 mm^3^, or until receiving a month of treatment. Time to humane endpoint was plotted as percent survival. ****P<0.0001, log-rank test. Middle, representative images are shown, scale bar = 1cm. Right, doubling times during treatment are reported. P = 0.0646, unpaired t-test.

The isogenic MCF10A panel presented here offers unique advantages and is complementary to current MYC-dependent models (Supplementary Figure 1F).^37–39^ Interestingly, the observed MYC-driven phenotype was exclusive to the *in vivo* xenograft assay. This highlights the importance of identifying a relevant context when studying MYC-dependent transformation, as our *in vitro* studies demonstrated little oncogenic MYC activity towards proliferation or *in vitro* acini transformation, and in no way was able to predict the striking *in vivo* response. Moreover, studying cancer biology using *in vivo* models most accurately reflects human disease and incorporates the contribution of tumour microenvironment and cellular stress. The *in vivo* context is therefore ideal for studying deregulated MYC and identifying MYC-dependent vulnerabilities as the relevant stress-signaling pathways will be active.

We have demonstrated that this model can recapitulate human BC at both the molecular and pathological level, and is transformed by deregulated MYC. We anticipate that this model will be a useful tool for identifying the context-dependent functions and vulnerabilities of MYC-driven cancers, that are more difficult to discover using traditional *in vitro* or murine models. Additionally, this system is reproducible and scalable; ideal characteristics for the identification and validation of anti-MYC therapies. Indeed, we have recently demonstrated the utility of this model through the validation of the G9a histone methyltransferase as an epigenetic, therapeutic target in MYC-driven cancers.^40^ We anticipate that this system will be able to predict the efficacy of novel therapeutic drugs targeting MYC and will identify novel vulnerabilities of MYC-dependent BCs.

## METHODS

### Cell culture

MCF10A epithelial cells were a kind gift of Dr. Senthil Muthuswamy and were cultured in MCF10A culture media, as previously described.^31^ Cell culture images were taken with a 32X(Ph) objective on an AxioObserver microscope.

### Lentiviral vectors

cMYC and PIK3CA^H1047R^ (cloned from Addgene plasmid # 12524) alleles were first cloned into pENTR4 no ccDB (686-1) (Addgene plasmid # 17424) and subsequently shuttled into pLenti CMV/TO Puro DEST (670-1) (Addgene plasmid # 17293) and pLenti CMV Hygro DEST (w117-1) (Addgene plasmid # 17454) respectively.^41^ Tetracycline-inducible cells were generated using pLenti CMV TetR Blast (716-1) (Addgene plasmid # 17492).

### Antibodies

Immunoblotting was performed against pAKT^S473^ (Cat# 4051, Cell Signaling), total AKT (Cat# 4685, Cell Signaling), MYC (9e10, homemade), Actin (A2066, Sigma), E-Cadherin (Cat# 3195, Cell Signaling Technology), Vimentin (Cat# 5741, Cell Signaling Technology) and Tubulin (Cat# CP06, Millipore).

### EdU incorporation assay

The MCF10A isogenic panel was grown for 24 hours in fully supplemented media and allowed to reach 50% confluency. At 50% confluency, cells were treated with EdU for 4 hours, processed according to manufacturer’s instructions (Click-iT EdU Alexa Fluor 488 Flow Cytometry Assay Kit, Cat# C10420, Life Technologies) and analyzed for EdU positivity using the LSR II flow cytometer (BD Biosciences) and FlowJo v10 software.

### Three-dimensional Matrigel morphogenesis assay

Cells were cultured on Matrigel (Cat# 354230, Corning) and in specialized MCF10A media for three-dimensional cultures, for 12 days as previously described.^31^ Images were acquired on day 12 using a FLUAR 5x/0.25 NA lens on an AxioObserver microscope (Zeiss). Cell line groups were blinded and quantified based on qualitative assessment of acinar circularity and size. Small round acini were considered normal; large and disorganized acini were considered transformed. One-way ANOVA with Bonferroni post-test for multiple testing was used for analysis.

### Xenografts

1×10^6^ cells were suspended in 50% matrigel (Cat# 354262, Corning) to a final volume of 0.2 mL and injected subcutaneously into the flanks of female NOD-SCID mice. For Tetracycline repressed cell lines, 100 μg/mL DOX (Sigma) was supplemented into the drinking water, prepared fresh twice a week. One-way ANOVA with Bonferroni post-test for multiple testing was used for analysis.

### Immunohistochemistry

Extracted tumours were fixed in 10% buffered formalin (Sigma) for 24-48 hours at room temperature and then stored in 70% EtOH at 4°C. Samples were then paraffin embedded, sectioned at 4 μm thickness and stained with hematoxylin and eosin. Ki67 (NB110-90592, Novus), TUNEL (homemade by PRP), ER (ab80922, Abcam), PR (NCL-PR312, Leica), HER2 (RM9103, Thermo Scientific), CK5 (NCL-LCK5, Leica), EGFR (28-0005, Invitrogen), p63 (VP-P960, Vector) and SMA (M0851, Dako) were used to further analyze tumour sections. All immunohistochemistry was performed by the Pathology Research Program Laboratory (PRP, University Health Network, Toronto, Canada).

### RNA sequencing analysis

Tumours were extracted and immediately flash frozen. RNA from xenograft tissue was harvested using the RNeasy Plus Universal Kit (Cat# 73404, Qiagen) following the manufacturer’s protocol. RNA was sent to the Princess Margaret Genomics Centre and sequenced using the Illumina NextSeq 500 for an average of 40M reads per xenograft growth. Fastq files were filtered using Xenome in order to filter out mouse contamination. Only reads mapping exclusively to the mouse reference were filtered out. The remaining reads were mapped to the human reference genome (hg19) and read counts estimated using STAR. Expressed genes were defined as requiring at least 10 reads in at least 2 samples. Read counts were upper quantile normalized. Finally, EBSeq was used to test for differential expression between experimental conditions. Genes were determined to be differentially expressed given an FDR threshold of 0.05.

### Gene ontology and gene set enrichment analysis

Differentially expressed genes between 10A.PE and 10A.PM were first ranked (rank = sin(postFC)*(-LOG(PPEE)) and then used for GSEA with GO Biological Process gene sets. Significantly up-regulated or down-regulated GO processes were then used to generate a cytoscape map. We acknowledge our use of the gene set enrichment analysis, GSEA software, and Molecular Signature Database (MSigDB).

## ACKNOWLEDGEMENTS

We would like to thank the Animal Resources Centre, Pathology Research Program Laboratory and Princess Margaret Genomics Centre teams for their advice, support, and expertise.

This work was supported with funding from the Tier 1 Canada Research Chairs Program (LZP), Canadian Institutes of Health Research (LZP).

---

**Supplementary Table S1 – MYC driven 10A.PM xenograft differential expression analysis.** Differential expressed genes from three 10A.PM and 10A.PE xenograft tumours. Only significantly changing genes that have a Log2 fold change of greather than 2 are shown. Genes with a positive fold change are increased in 10A.PM relative to 10A.PE. Genes with a negative fold change are decreased in 10A.PM relative to 10A.PE. realFC: real fold change, postFC: posterior fold change, PPEE: posterior probability of being equally expressed, PPDE: posterior probability of being differentially expressed.

**Supplementary Table S2 – Summary of GO-term gene set enrichment analysis.** Differentially expressed genes in 10A.PM tumours were analyzed by GSEA using GO-term gene sets. Grouped nodes in Figure 3B contain the GO-terms listed here. A summary of all GO terms analyzed are also included.

**Supplementary Table S3 – Summary of MYC and Breast cancer GSEA results.** Differentially expressed genes in 10A.PM tumours were analyzed by GSEA using MYC and breast cancer gene sets. NES: normalized enrichment score.

**Supplementary Figure S1.**
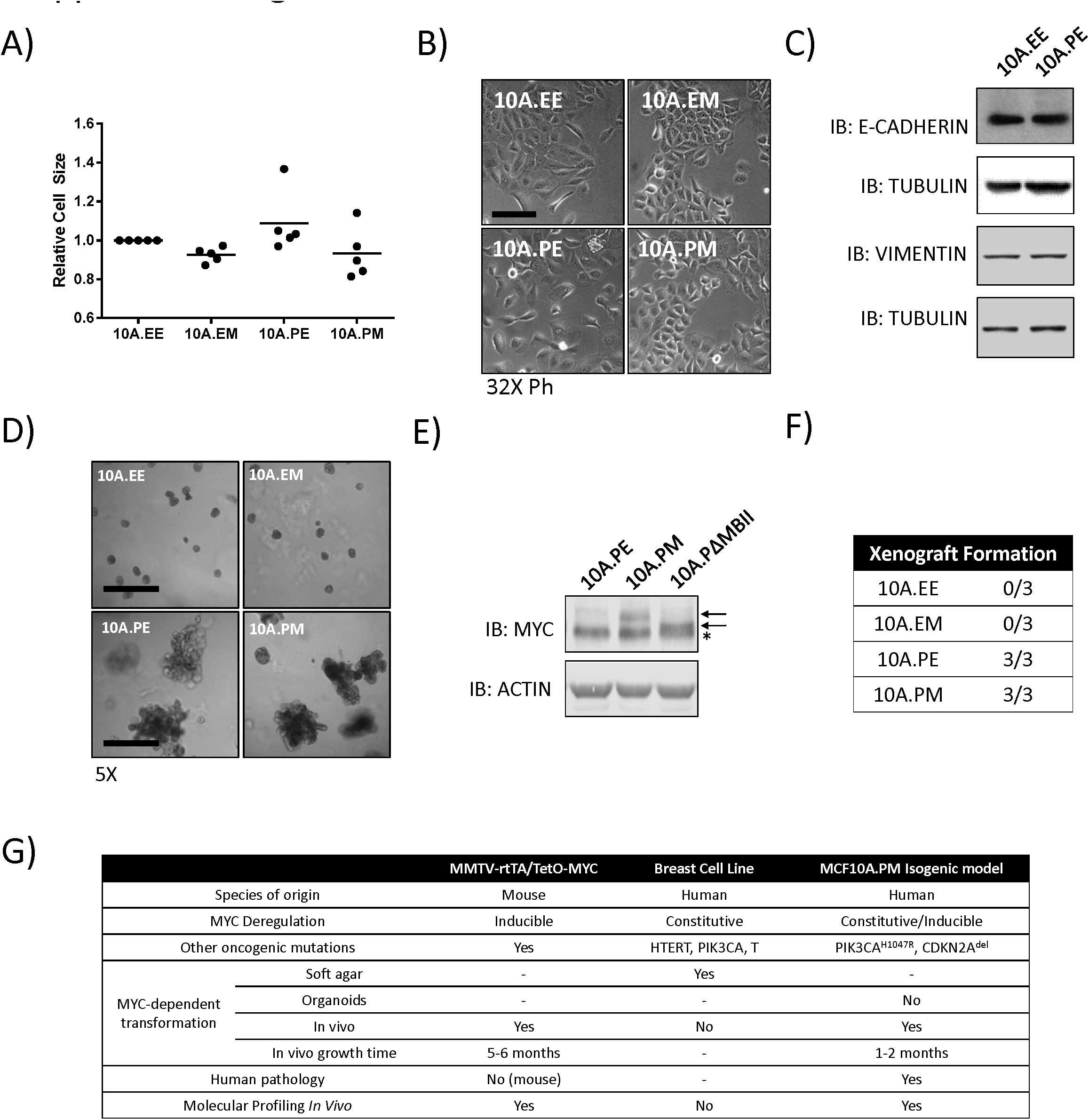
**(A)** Flow cytometry measurements for the isogenic MCF10A panel. Mean values are shown for relative cell-size, one-way ANOVA with Bonferroni post-test for multiple testing, 5 biological replicates. **(B)** The MCF10A isogenic panel was cultured in standard MCF10A media for 48 hours before imaging to monitor cell morphology. **(C)** 10A.EE and 10A.PE were cultured for 48 hours and protein lysates were harvested to monitor markers of EMT. **(D)** Images were acquired on day 12 and quantified based on assessment of acinar circularity and size. Scale bar = 500 μm **(E)** Ectopic MYC expression from cells injected into NOD-SCID mice. Arrows point to ectopic WT and ΔMBII protein. * indicates endogenous MYC. **(F)** The MCF10A isogenic panel was injected into female NOD-SCID mice as previously described. Summary of tumour formation is presented, 3 animals per group. **(G)** Summary table of common GEMM and breast cell line models in comparison to the MCF10A.PM isogenic model. “-” represents not tested.

